# Electrostatically Assembled Layer-by-Layer Plasmonic Implant for Combination Therapy against Multidrug-Resistant Chronic Wound Biofilms

**DOI:** 10.64898/2026.05.29.728676

**Authors:** Nutan Shukla, Ratnesh Das

**Author notes:** Corresponding: Nutan Shukla.

## Abstract

Although persistent infection from chronic wound lowers the efficacy of single therapy; But combination therapy with prolonged drug release have shown promising effect, such as elimination of heavy bacterial film and multi/single drug resistance with minimal side effects. One such breakthrough is multilayered scaffold that are chemically and physiologically stable. To achieve this objective, we demonstrated construction of layer-by-layer (LBL) assembly aligned by alternate deposition of (PEI/PSS and PAA/PSS) based on electrostatic force. These films exhibited loading of Ibu and gen which can be altered depending on different parameter such dipping time, pH and number of layers. Briefly, gen was encapsulated onto PPGNRs, as GNRs have good stability and their antibacterial properties can be enhanced by tunning surface upon adding chemical drugs. Such functionalized GNRs showed excellent photo thermal property which assisted controlled release of gen via deconstruction of thin layers, whereas burst release of ibu. Furthermore, complete disruption of bacterial colonies when combined with near infrared irradiation (NIR). The formed LBL endowed the great healing capability, controlled antibiotic release solve the problems of bacterial resistance due to synergistic effect. Taken together, the antibacterial, cytocompatibility, and stimuli responsive characteristics of this robust multilayer assembly can be promising multifunctional drug delivery system in different medical aliments.

## Introduction

Chronic wounds remain a major clinical challenge because they fail to progress through the normal phases of healing, including hemostasis, inflammation, proliferation, and tissue remodeling [1,2]. Such wounds are commonly associated with diabetes, vascular insufficiency, pressure injury, post-surgical complications, and persistent microbial contamination. Unlike acute wounds, chronic wounds usually exhibit prolonged inflammation, high exudate production, impaired angiogenesis, excessive protease activity, and delayed re-epithelialization [1]. These pathological features create a wound microenvironment that is highly favorable for bacterial colonization and biofilm formation, ultimately delaying tissue repair and increasing the risk of severe infection [2,3].

One of the most serious obstacles in chronic wound treatment is the formation of bacterial biofilms. Biofilms are organized microbial communities embedded within a self-produced extracellular polymeric substance matrix, which protects bacteria from host immune responses and reduces the penetration and efficacy of conventional antibiotics [3,4]. In biofilm-associated infections, bacteria may show substantially higher tolerance to antimicrobial agents than planktonic bacteria, often requiring repeated or high-dose antibiotic administration [4,5]. This prolonged antibiotic exposure contributes to the emergence of multidrug-resistant pathogens and further complicates wound healing. In chronic wounds, pathogens such as *Staphylococcus aureus*, methicillin-resistant *S. aureus, Pseudomonas aeruginosa*, and other Gram-negative bacteria can persist within biofilms, causing recurrent infection, inflammation, tissue necrosis, and delayed closure [3,5]. Therefore, therapeutic approaches that combine antibacterial activity with biofilm disruption are urgently needed.

Conventional wound dressings mainly provide a passive barrier and moisture control; however, they are often insufficient for complex infected wounds. In contrast, advanced drug-delivery wound dressings can actively release therapeutic molecules in response to wound conditions [1,6]. Sequential or combination release of multiple therapeutic agents is especially attractive because chronic wounds require simultaneous control of pain, inflammation, bacterial infection, and tissue regeneration. For example, incorporation of anti-inflammatory or analgesic agents such as ibuprofen can help reduce local pain and inflammation, while antibiotics such as gentamicin can suppress bacterial growth. Ibuprofen-releasing foam dressings have been clinically investigated for painful exuding wounds and have shown relevant pain-relieving effects in chronic wound patients [7,8].

The pH of the wound microenvironment is another important factor in wound progression and infection. Healthy skin is generally mildly acidic, whereas chronic and infected wounds often shift toward neutral or alkaline pH [9]. Elevated wound pH has been associated with higher bacterial burden, impaired oxygen release, altered protease activity, and delayed healing [9,10]. This pH variation provides an opportunity to design smart wound dressings that respond to wound exudate and pH changes. Materials capable of pH-responsive swelling, degradation, or drug release are therefore highly useful for chronic wound applications, particularly when combined with antimicrobial strategies.

Layer-by-layer assembly is a versatile technique for fabricating thin, multifunctional films through the sequential deposition of oppositely charged polyelectrolytes, nanoparticles, proteins, or therapeutic molecules [11,12]. The driving forces for LBL film formation include electrostatic interaction, hydrogen bonding, hydrophobic interaction, and specific molecular recognition. This method allows precise control over film thickness, surface charge, roughness, wettability, drug loading, and release behavior by changing the number of bilayers, pH, ionic strength, polymer composition, and dipping time [11–13]. Because of these tunable properties, LBL films have been widely investigated for biomedical coatings, antimicrobial surfaces, tissue engineering, and controlled drug delivery [12,13].

Polyelectrolyte multilayers are particularly suitable for wound dressing applications because they can host both hydrophilic and hydrophobic therapeutic agents within different regions of the film [11,13]. Positively charged polymers such as polyethyleneimine can provide strong electrostatic interaction with negatively charged polymers such as poly (sodium 4-styrenesulfonate) or poly(acrylic acid), enabling stable film growth. At the same time, polymeric domains within the multilayer can act as reservoirs for drug molecules, allowing either rapid release or prolonged release depending on drug–polymer interaction strength and environmental conditions. In the present system, the LBL structure is designed to provide a stable multilayer scaffold, where ibuprofen is incorporated for early burst release and gentamicin-loaded gold nanorods are embedded for NIR-responsive antibacterial therapy.

Photothermal therapy has emerged as a promising non-antibiotic antibacterial strategy for infected wounds. Under near-infrared irradiation, photothermal agents absorb light and convert it into localized heat, which can damage bacterial membranes, disrupt biofilm structure, and increase bacterial sensitivity to antibiotics [14–16]. Compared with systemic antibiotic treatment, NIR-triggered photothermal therapy offers spatial and temporal control, deeper tissue penetration, and reduced systemic toxicity. Importantly, because photothermal killing is based on physical membrane damage and protein denaturation rather than a single biochemical target, it may reduce the likelihood of conventional drug resistance [14,15].

Gold nanorods are among the most widely investigated photothermal nanomaterials because their longitudinal surface plasmon resonance can be tuned into the near-infrared region by controlling their aspect ratio [14,16]. Upon NIR irradiation, gold nanorods generate localized heat with high photothermal efficiency. Their anisotropic shape, optical tunability, and surface modification potential make them attractive for antibacterial wound therapy. However, direct use of bare gold nanorods can be limited by colloidal instability, surfactant-associated toxicity, aggregation, and uncontrolled interaction with biological components. Therefore, polymer coating or encapsulation is often used to improve stability, reduce toxicity, enhance dispersibility, and enable drug loading [14,16].

Combining gold nanorod-mediated photothermal therapy with antibiotic delivery is a rational strategy for eradicating biofilm-associated infection. Localized heating can increase bacterial membrane permeability and disrupt the extracellular polymeric matrix, allowing improved antibiotic penetration into the biofilm [14–17]. At the same time, antibiotic release at the wound site can reduce the required therapeutic dose and minimize systemic exposure. This synergistic combination is especially valuable for multidrug-resistant infections, where single-mode therapy often fails. Recent studies have shown that gold nanorods and other photothermal nanomaterials can enhance antibacterial activity under NIR irradiation and promote wound healing when integrated into suitable biomaterial platforms [15–17].

Despite these advances, several limitations remain in the development of smart antibacterial wound dressings. Many systems rely on single-drug release, lack sequential therapeutic control, or fail to combine pain/inflammation management with biofilm eradication. Other systems show strong antibacterial activity but may have limited biocompatibility, poor structural stability, or uncontrolled burst antibiotic release. Therefore, an ideal wound patch should provide: first, a stable and tunable scaffold; second, early release of an anti-inflammatory or analgesic agent; third, controlled antibiotic delivery; fourth, externally triggered biofilm disruption; and fifth, acceptable cytocompatibility with normal skin-associated cells.

In this study, a multifunctional LBL wound patch was developed using electrostatically assembled polyelectrolyte multilayers for combination therapy against chronic wound-associated bacterial biofilms. The system integrates ibuprofen for early pH/exudate-assisted burst release and gentamicin-loaded polymer-coated gold nanorods for NIR-triggered controlled antibacterial release. Upon NIR irradiation, the gold nanorods generate localized heat, promoting photothermal disruption of bacterial biofilms and controlled gentamicin release. This dual-action strategy is expected to reduce inflammation and pain during the early stage while providing enhanced antibacterial activity during the controlled release phase. The designed LBL nanoscaffold therefore represents a promising smart wound patch for MDR biofilm-associated chronic wound therapy.

## Material and methods

a. **Chemicals-**Buffer (ph. 4, 5 and 7.4), Linear polyethyleneimine (LPEI, Mw=25000), Poly (sodium 4-styrene-sulfonate) PSS, Gentamicin sulfate (GS)
b. **Equipment:** To conduct NIR release experiment infrared camera (Fluke, USA) was used. H^1^NMR was used to analyze the chemical synthesis ECX-400 instrument (JEOL, Japan). To characterize, the TEM image was obtained via JEOL 1000CX (JEOL, Japan) operating at 80keV and the modified GNRs were monitored using a Nicolet iS 10 FT-IR spectrometer (Thermo Scientific, USA). The charge density was acquired using Zeta potential analysis by Dynamic Light Scattering (DLS) Zetasizer. The spectral absorption was quantified by Neosyss-2000 UV-vis spectrophotometer (SCINCO, USA) and Fluorescent spectrometer QM-400 (Horiba Scientific, USA). The cell counting and viability was measured via micro plate reader (Tecan infinite series 200, Germany). The cell sorting after staining was obtained by (fluorescence-activated cell sorting (FACs) caliber, BD Bioscience).
c. **Methods**
  1. **Preparation of PPGSTGNRs (Drug 2)** the formation of PPGenGNRs was obtained using electrostatic interaction. The interaction between TGNR and PPEG generates hydrophobic pockets which suitable for encapsulation of different types of drugs, which is suitable to maintain the actual effect of pharmaceuticals. Briefly, Aqueous suspension of TGNRs (20.3×10^-9^M, 3.5 mL) in D.I was introduced drop wise into (polymer drug) solution PPEG: GS aqueous suspension (20mg, 10ml, 5mg) resulting mixture was covered and allowed for stirring for 48h at temperature 30^0^ C. Over the time to obtained purified form of PPGSGNRs, it was applied for centrifugation at 10000 RPM for 20min; pellet was collected and supernatant was removed. To check stability the PPGSGNRs was lyophilized and later it was dispersed in different solvent (PBS, D.I) and peak absorption with LSPR and TSPR shift was confirmed by Uv-vis spectrophotometer. Since, GS do not have strong Uv-vis absorption, the encapsulation amount of GS was determined via HPLC peak.
  2. **Assembling of multilayered film of polyelectrolyte LPEI/PSS (Base layer/Sample 1)** The PE multilayer film was prepared using alternate assembly of LPEI/PSS by dipping in the PE solutions. The driving force between PE helped their adsorptions sequentially. In brief, silica wafer or substrate was activated by exposing to oxygen plasma (OS) for 10min and cleaned with EtoH.The adsorption of PE was employed by continuous dipping for 5min and washing for 1min in each PE solution for example-(LPEI/PSS), this cycle was repeated until to obtain required number of layer on silica wafer. In this research the LPEI is used initially to coat bare silica to maintain the strong assembly of further layers. The sample with different layer/thickness was prepared by optimizing dipping and washing time. Before, starting for drug layer, the above sample was dried well to avoid aggregation of multiple layers

To prepare drug coating layer, the primary study was performed, the interaction between model drug Rhodamine 6 B and PSS was evaluated in different pH (3-10). This study can be useful to load different types of drug based on affinity towards different types of polyions.The binding of RB to PSS is depended on protonation degree in different pH, this phenomenon can be useful to load drug in the hydrophobic domain of polymer domain. (10.1021**/jp061457j CCC**). Later the RB was replaced with anti-inflammatory hydrophobic drug namely IBU, following previous method the association of IBU into the hydrophobic domain of polymer was obtained by dipping the **sample 1** into the polymer drug solution (IBU: PSS, 5mg;20mg in D.I) until the required amount of IBU was loaded named as (Drug 1)(https://doi.org/10.1002/1439-2054(20011001)286:10<591::AID-MAME591>3.0.CO;2-I)

To prepare Drug 2 layer, same method as above was followed (refer to sample 1), briefly the obtained dried PPGSGNR was dispersed in D.I completely. The prepared sample 1 was dipped into the solution of PPGSGNRs until the effective loading was obtained as (LPEI/PPGSGNRs/LPEI/PSS). Once both (Drug1 and Drug 2) was loaded, the thickness at particular layer was obtained by profilometer and hydrophicity/hydrophilic was analysis by change in contact angle.

### Evaluation of thermal elevation efficiency of PPGenGNRs

The near infrared (NIR) light was studied by irradiating sample surface. The increase in temperature was monitored using infrared camera at two different laser power (0.2 and 1.6) and blank silica substrates without multilayer for comparative studies. To demonstrate the thermal expansion responding to NIR, the prepared substrate (sample1 and drug 2) non-coated silica wafer (**control 1**)and only pillion coated silica wafer(**control 2**)was exposed to NIR with different laser power ranging (0.2, 0.4, 0.6, 0.8, and 1.6 W/cm^2^) and NIR triggered PTT conversion ability and elevation of Temperature was monitored using infrared camera. The change in temperature was noted at particular time and total irradiation time was for 15min. Temperature changes and photostability of this conjugate were clarified by repeating 3 times to minimize possible errors and results were noted.

### 2.6. Release studies of a) IBU b) GS

To evaluate the release of IBU, the prepared LBL with loaded (IBU+ polymer) was dipped into different pH (5 and 7.4) at 30^0^ C and different time point. The obtained IBU release at different pH was subjected to HPLC and IBU peak was observed, simultaneously the temperature stimuli release of GS was obtained at different irradiation times, **PPGSGNRs** LBL was placed at different temperature (R.T, 37^0^ C and 65^0^ C) in PBS for 120 min. Later, 0.5ml was removed and replaced with fresh PBS with same volume at define time point. The samples were centrifuged at 13000 rpm for 15 min and the amount released of free GS was measured by HPLC

### Biocompatibility assessment using live and dead staining

To assess the biocompatibility, the NIH3T3 cells were used to demonstrate the effect of our Nano scaffold in normal cell environment. Briefly, the 5×10^3^ Cell/well was seeded in 12 well and incubated for 24h in culture medium. After 24h, the prepared LBL with loaded drug and without loaded and immersed with cells and examined at different time (0, 4h, 12h and 24h). The culture medium was changed over the set time point and replace with fresh medium. The cells morphology was confirmed by optical microscopy, whereas viability was confirmed via staining with PI and FDA following previous protocol. Additionally, **Migration assay** was performed in which the 5×10^3^ cells were seeded in 12 well and scratch was provided using sterile tip. The cell debris were washed using fresh media, the LBL scaffold with/without film was placed on the scratch and irradiated with NIR and allowed for incubation at different time. After incubation, the cells were stained and observed under the microscope.

### Antimicrobial susceptibility study of release of GS from films

The minimum inhibitory concentration (MIC) of GS was obtained from previously demonstrated studies. To evaluate the combination effect we used well diffusion method, agar was punched of suitable diameter into the prepared Muller Hinton agar plate (MH).With the help of microtip the obtained gram positive and gram negative pathogens were inoculated onto the MH agar plate and incubated for 24h. Over the time, known PPGenGNRs concentration was added into the agar well in each of pathogen and irradiated with different laser power (0.2 and 1.6 W/cm^-2^) for 1, 5 and 10 min, incubated for 24 h. Over the time zone of inhibition of test and control sample was obtained by measuring diameter in millimeters.

## Discussion and results

**Figure.1**. illustrates the assembly of polyelectrolyte via layer by layer (LBL) which was deposited on pretreated silicon wafer. ^**[39-42]**^ The prepared nanoscaffold displayed several properties like 1) Dual drug loading (pain killer and anti-inflammatory hydrophobic ibuprofen and hydrophilic antibiotic Gentamicin sulfate) 2) The burst release of ibuprofen can be controlled in different pH due to its solubility and wound exudate 3) The assembled layer enabled the loading of GNRs encapsulated gentamicin and released upon responding to external stimuli.^**[32-38]**^It is informed that over 70% of wound infection acquired resistance due to continuous overdosing in chronic wound resulted from Diabetes, post-surgery and so on, therefore investigation of novel approach / therapeutic is in need to tackle chronic wound environment^**[43]**^ Our system demonstrated the successful entrapment of GNRs which was pre-loaded with Gen into amphiphilic network of PP to elongate circulation time.^**[44]**^ As many research demonstrated the efficiency of plasmonic nanomaterial’s namely GNP, AgNP or GNRs for combination therapy, which is effective over single therapies. Beside all, GNRs with extreme surface parameter i.e. strong LSPR and TSPR in the range of NIR (600-1100nm), fine tuning has become ideal candidate for PTT based treatments.^**[45]**^In presence of GNRs when wound is irradiated with NIR the perfusion rate is enhance and bacterial cell wall is disrupted due to localized heat this leads to continuous release of wound factors like heat shock proteins (HSPs) and immune cells which is needed for remodeling, regeneration and homeostasis of surrounding tissues, thereby speedup healing time. ^**[46-50]**^

**Figure 1.**
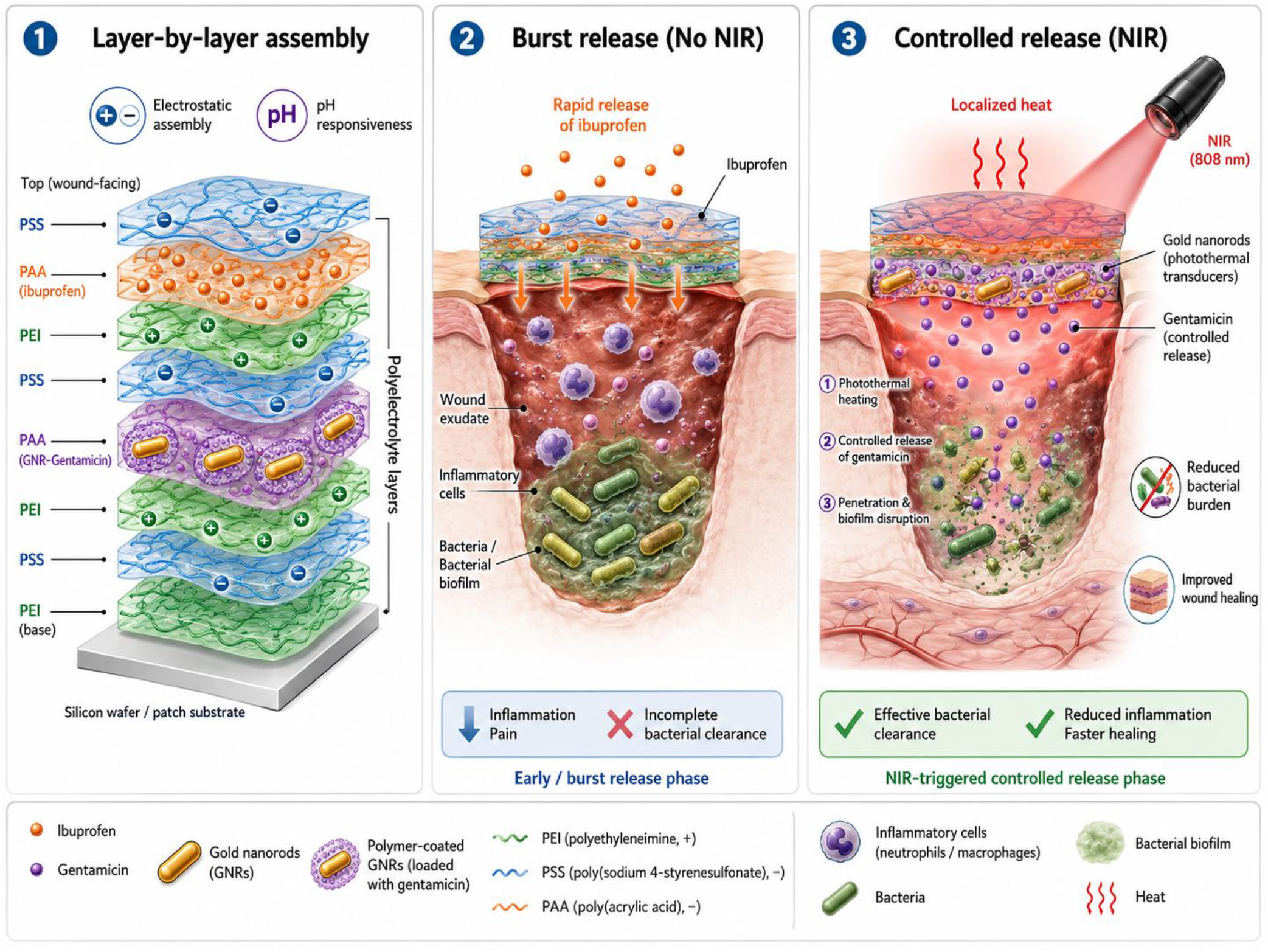
Schematic illustration of smart wound patch: **a)** Shows LBL assembly and dual drug loading and release. **b)** Burst and controlled drug release at wound site. **c)** Synergistic disruption of bacterial film responding to NIR.

The fabrication of CGNRs followed via seed mediated method enables surface functionalization using pre-synthesized thiol moiety” (SH). This not only increased surface charge, but enhanced solubility and stability in different solvent for several months, the prepared CGNRs and TGNRs were characterized by spectral shift obtained using Uv-vis spectrometer. The red shift from 730nm to 780-nm this displayed the successful formation of TGNRs and a Zeta potential result demonstrated the change in surface charge from 21mV to 41mV **(Fig.1B)**. ^**[51]**^ Such functionalization approach is essential for further conjugation of different molecules such as (drug or polymer). ^**[52]**^ The chemical modification of therapeutic using different covalent bond have reduced the inherent effect, thus the physical encapsulation of drug into the hydrophobic pocket of amphiphilic polymer has become ideal approach for prolonged stability. ^**[53]**^The strong cationic charge from TGNRs facilitated electrostatic adsorption of amphiphilic PP **(Fig.1A)** The Uv-vis and zeta potential results displayed spectral shift from (40mV to15mV) **(Fig.1B)**. The spectral broadening in the longitudinal were observed, this characteristic is important for further NIR studies. In addition, morphology of CGNRs, TGNRs and PPGNRs was confirmed by TEM **(Fig.1C)**. ^**[54]**^

**Figure 1.**
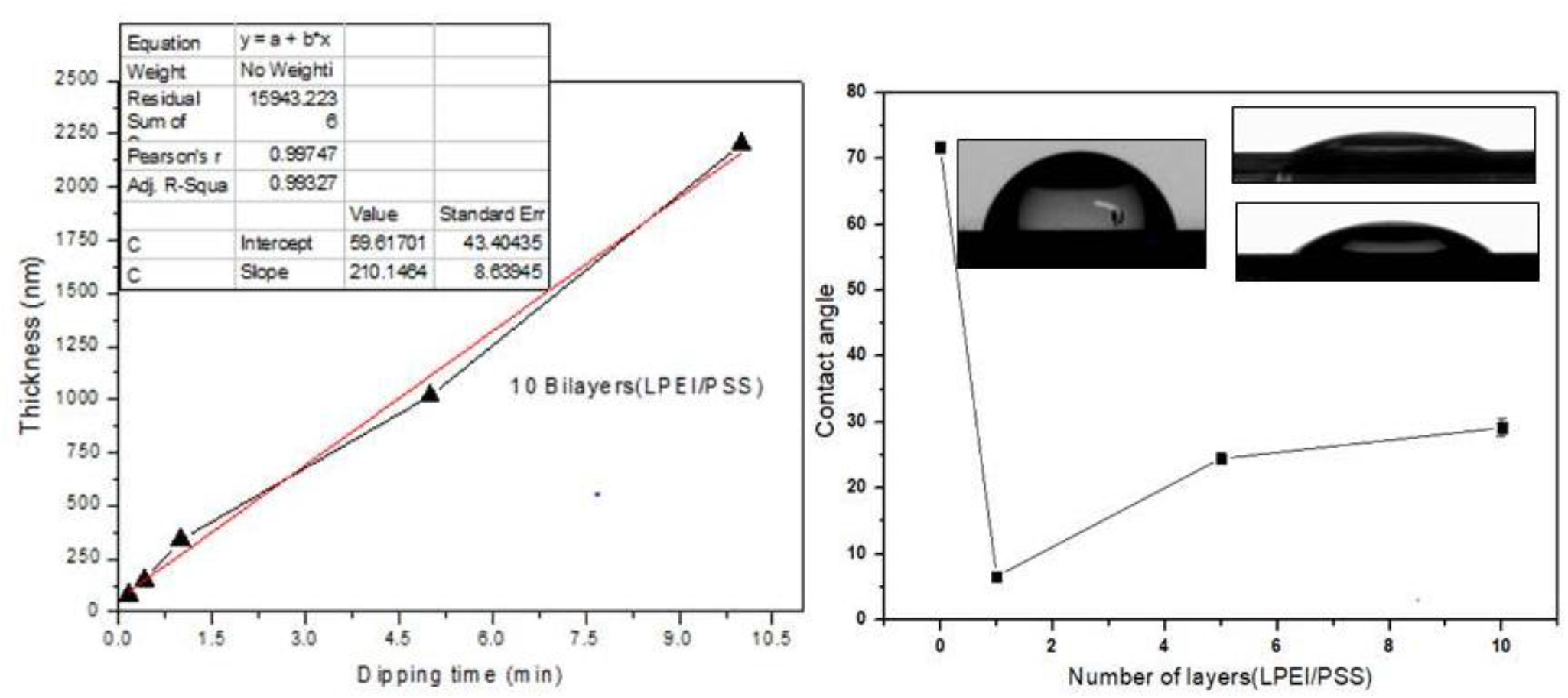
Thickness and Contact angle before and after LBL assembly **a)** zero layer **b)** 1bilayer **c)** 5bilayer **d)** 10 bilayer-(No drug layer)

To encapsulate antibiotic GS was formulated with (1:10 ratios of GS: PP) and dried as film. ^**[56]**^ TGNRs was added slowly, since GS does not show Uv-vis for loading efficiency the full release was analyzed using HPLC.^**[55]**^ As combination of nanoparticle drug is cyclically absorbed throughout the film development process which is one of the structural components observed growth behavior of LbL.^**[57]**^The thickness of film can be alter depending on medical application such systems have been the subject of several research.^**[58]**^When a single stone-diffusive component absorbs into the majority of a multilayer film that is developing, exponential growth is typically seen, and each consecutive cycle of deposition results in increasing amounts of material absorbed. ^**[59]**^ To determine successful formation of LBL with (GS and IBU), the film thickness, and roughness and contact angle was measured after each cycle of dipping or washing shown in **Figure (2c)**. ^**[60]**^ Film thickness is important parameter as this decides the overall arrange of LBL, when referring **Figure2 (a, b)** film thickness is increasing when the dipping time is increased; pH and ions influence the pattern of deposition of films. The over dipping disrupts the film formation and results into aggregation.

**Figure 2.**
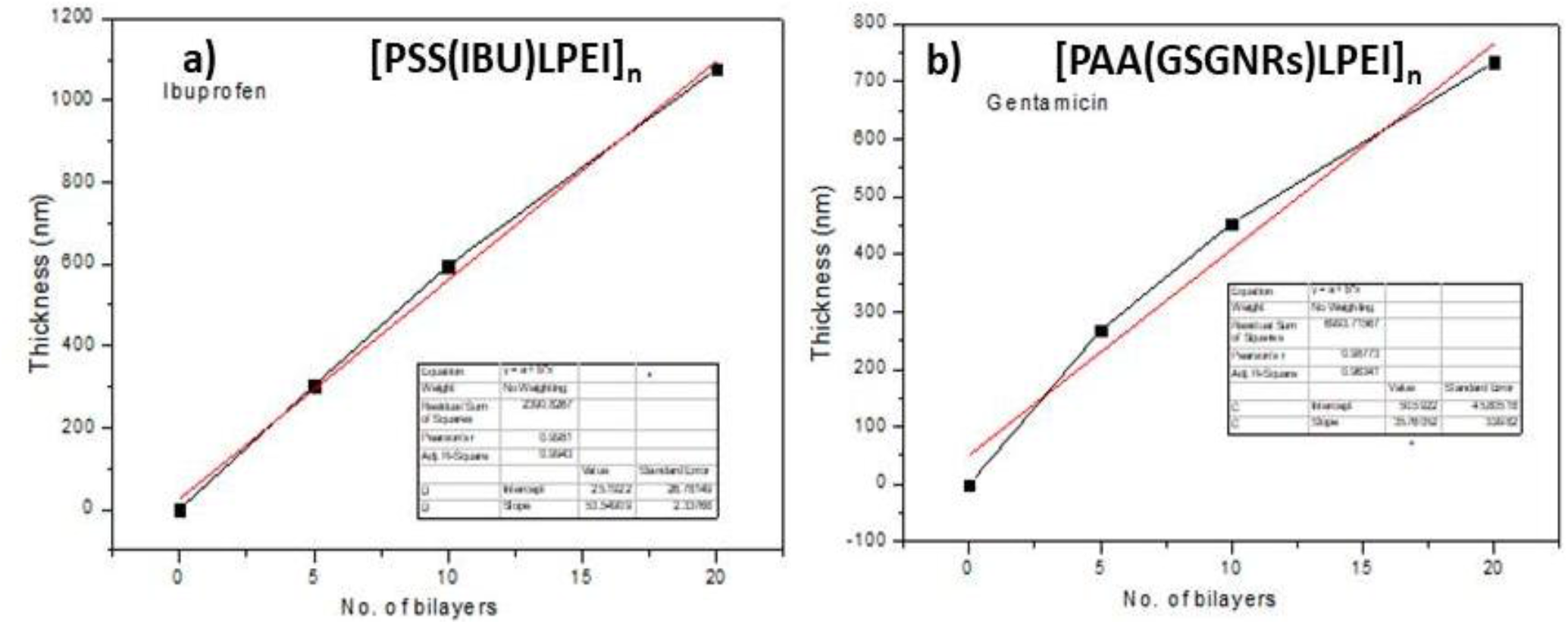
Drug loading **a)** Drug 1 (IBU) in PSS/LPEI **b)** Drug 2 GSGNRs in PAA/LPEI with increasing thickness upon increasing bilayers.

There are three basic material requirements that must be taken into consideration when designing layer-by-layer systems for the release of antibiotics in a physiological environment: (1) incorporation of a therapeutic, (2) a mechanism of release, and (3) any other materials required to permit stable layer-by-layer film growth without compromising biocompatibility. So, different biopolymers such as hyaluronic acid (HA), Chitosan, PAA have been reported for the delivery of gentamicin. ^**[61]**^The protonation degree in pH (4-5.5), the charge density difference in backbone of (PAA > HA), and molecular weight helps enables easy and high incorporation of drug as compared to any other polymer Thus PAA was used in our study for encapsulation PPGSGNRs. After each cycle of deposition PPGSGNRs/PAA, the protective layer of LPEI was deposited following dipping and washing which results into (LPEI/PPGSGNRs/PAA/LPEI) _10_. ^**[62]**^ The film thickness is monitored, and can be changed by changing different parameter such as pH, dipping time shown in **figure (2a)**, the growing film thickness change the surface quality **Figure (2c)**

Since our nanoscaffold enabled the multiple drugs loading option, our second drug was anti-inflammatory, pain reliever” IBU. Briefly to prove the working concept we conducted primary study in which we selected the model drug “RhoB6G” to load in the network of PSS which is already reported. Owing the protonation degree in different pH the RhoB6G is intercalated strongly inside the PSS. After successful encapsulation of RHoB6G, release rate was observed in different pH (5-10) Shown in **Figure (3a)**.

**Figure 3.**
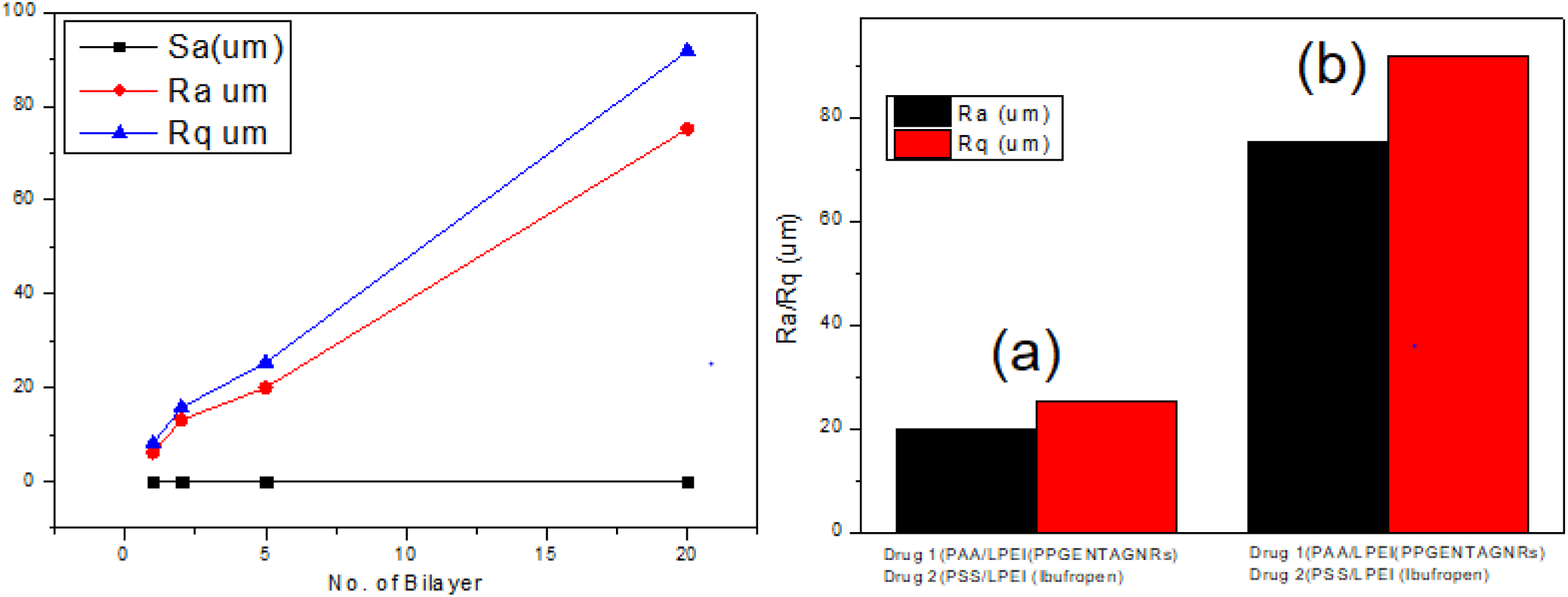
Characterization of using AFM-surface roughness analyses for increasing thickness of drug (1 and 2) layer depending on Number of bilayer

**Figure 3A.**
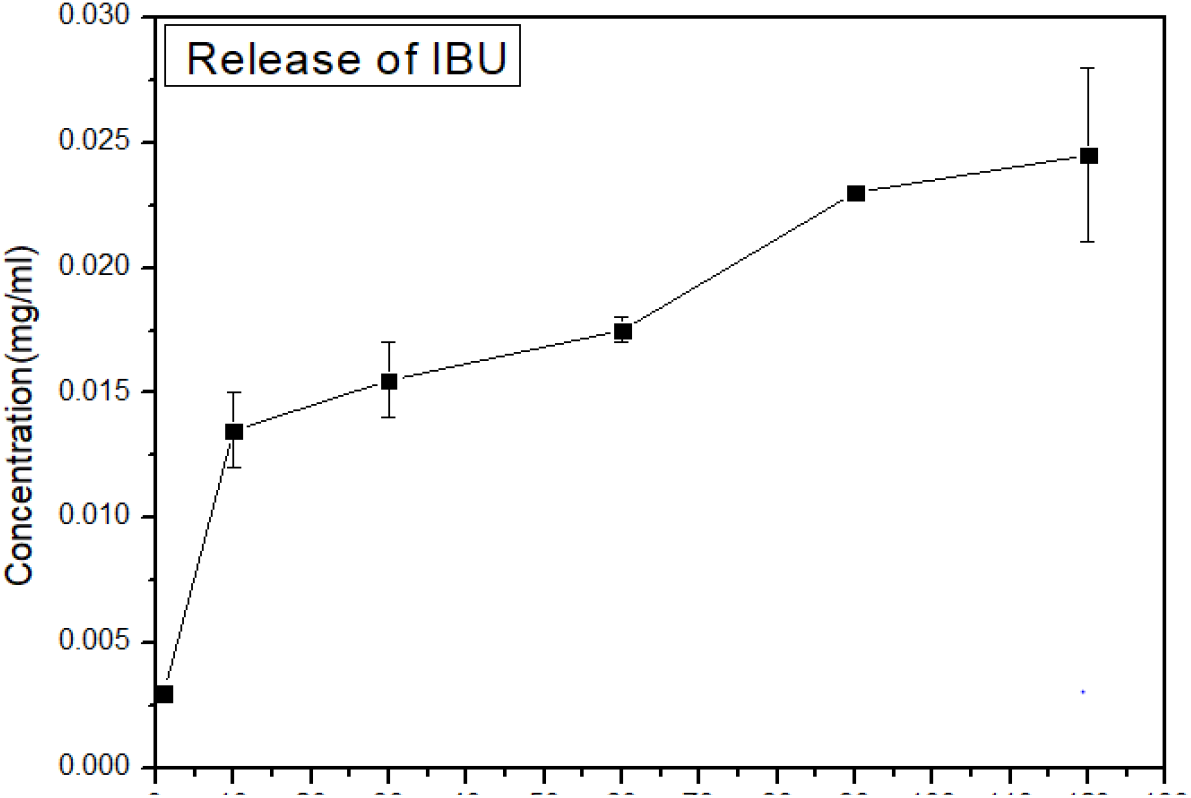
Burst release of IBU from PSS/LPEI

In pH 7,9 or 10 the release rate was high as compared to pH less than 5or 6 as shown in **Figure(3b)** This study can be useful to release the drug in chronic wound exudate where pH is from (7-10). Similarly, we followed the encapsulation of extremely hydrophobic IBU into the cavities of LPEI/ PSS, the film cavity is much higher than the IBU particle size. **[63]** The loading of IBU was obtained at pH 7. **[64]** The growing layer can affect the loading of IBU hence we prepared the 10 bilayers of (LPEI/PSS-IBU-PSS/LPEI) _10_, burst release was obtained in first 10 min **Figure (3a)**, whereas GS was found to be controlled release responding to NIR (0.6, 1.6W/cm^-2^) the loading of GS in 10 bilayer was about 82.5% (1.645mg), full release was obtained until the 60min. **Figure (3b)**.

**Figure 3B.**
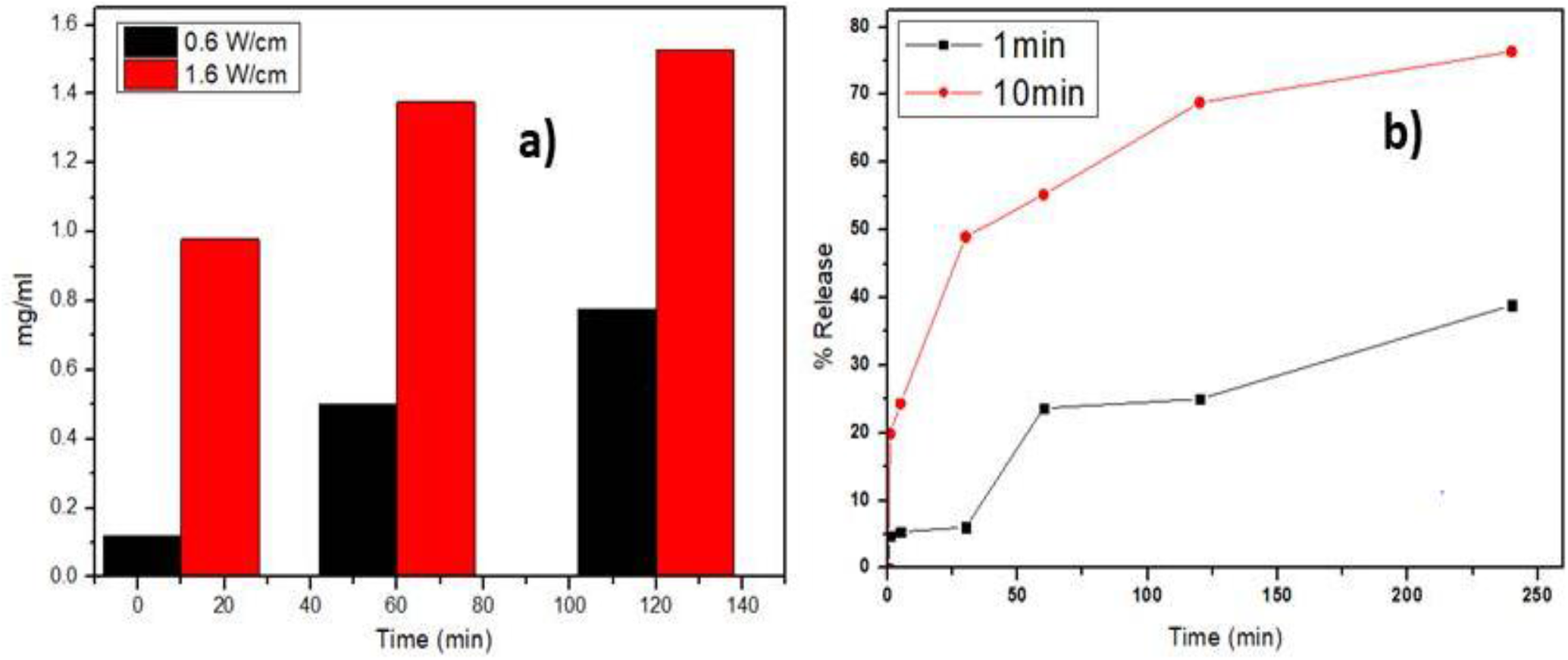
Controlled releases of GS from PPGNRs **a)** depending on laser power (0.6, 1.6W/cm^-2^) **b)** depending on time

The Cell adhesion and Cell differentiation was studied to demonstrate the biocompatibility of LBL film. Briefly the NIH3T3 was incubated with prepared film and incubated for 4h, 12h, 24h and 48h. After the incubation with respective time we provided the PI and FDA staining and viewed using CLSM shown in **Figure (4)**. The CLSM images revealed the morphology and lives cells. The FDA stain was retained in control (only silica substrate) and Test (LBL film) which explains that the prepared LBL is non-toxic even in 24h or 48h. **[65]** To check the proliferation rate or cell doubling, at the end of the experiment we trypsinized the cells and proceeded with trypan blue counting method, Although, there are different types of PE used in fabrication of LBL, it did not hampered the growth of cell this prove that this nanoscaffold is biocompatible. **[66]** Additionally, to study cell adhesion properties of the prepared LBL, NIH3T3 cell were allowed to attach onto the surface and incubated for 24h and optical viewed under the CLSM ref **Figure (4a)**.In wound healing one of the important points is the reepithilialization occurred when normal cells migrate and help in wound closure. The transfer of cells from one location to another occurs as a result of the multi-step, complicated process known as cell migration, which is crucial for wound healing. The wound scratch model is NIH3T3 test for assessing in vitro wound healing. After achieving 80% confluent of 3T3 cells that had been deprived of nutrients for 24 hours before being manually scratched to imitate a wound in order to test the ability of the prepared LBL to induce cell migration and proliferation **(Figure 6)**.

**Figure (4a).**
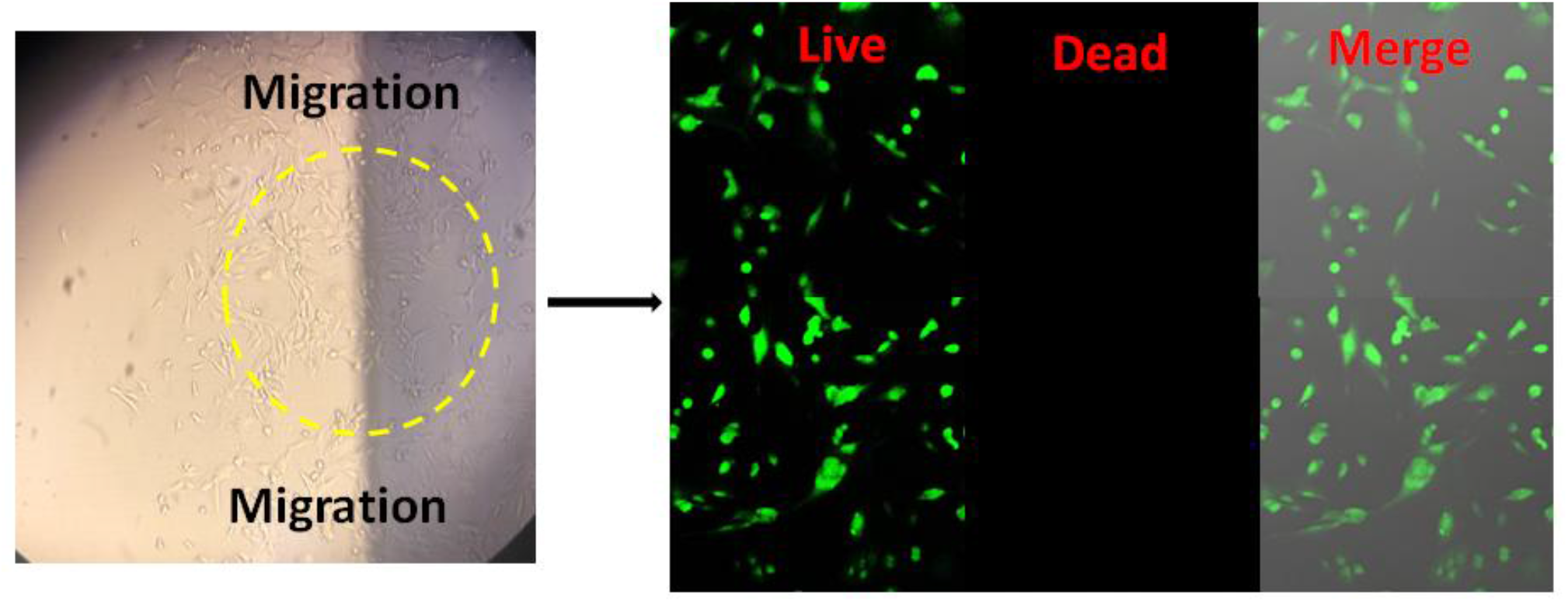
illustration of migration of NIH3T3 cells after scratch and incubating with LBL using live and dead staining

**Figure (4b).**
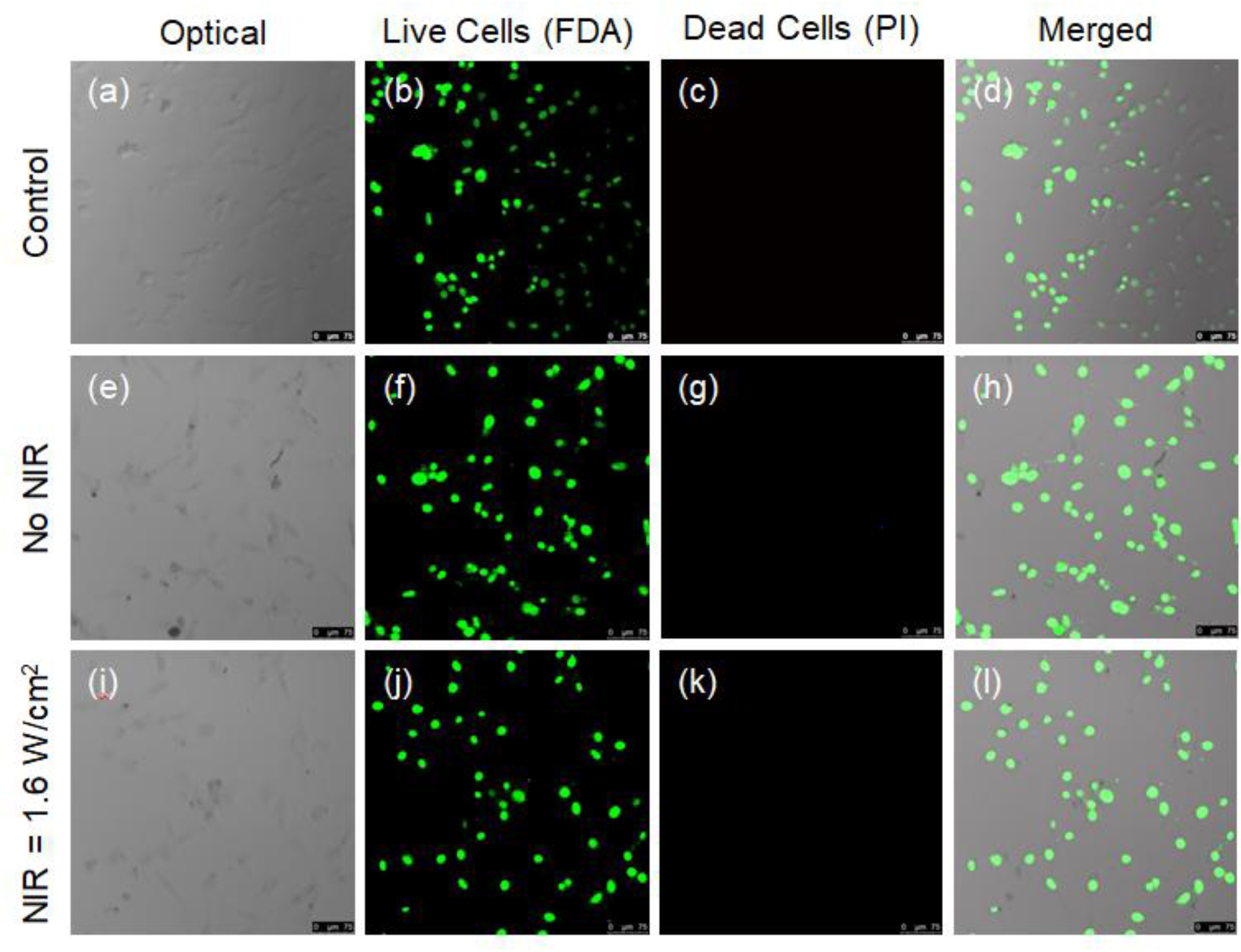
illustration of biocompatibility of prepare LBL using live and dead staining

Additionally, the growing interest in NIR due to its ability to release wounding healing marker at molecular level have marked them one of all best treatment option in present. However it is disrupted upon toxicity of prepared nanoplatform.To prove this concept we performed migration assay by incubating LBL in pre-scratched NIH3T3 cells and irradiated with different laser power (0.6, 1.6) for (1,5,10 min). The obtained microscopy explains the migration was improved in the NIR treated sample over the time.

Combining the investigation of alternative therapeutic option the combination of different modality has been illustrated continuously which has improve the etiology of chronic wound. Amongst, NIR have broad insights as an alternative tool to enhance treatment outcome. To further investigate the PTT effect, the prepared LBL was exposed NIR with different laser power and time, the heat/elevation in temperature was noted. The LBL consist of GNRs showed high increment in temperature upto 70^0^ C compared to control sample (LBL without GNRs) which showed minimal increase in temperature shown in **Figure (5)** The increase in total temperature is dependent on the number of layers and concentration of GNRs as reported in **chapter 1 and 2**. The exponential growth in film increases the encapsulation efficiency, when noting the temperature of LBL (without GNR) showed increase in temperature upto 20^0^ C.

**Figure 5.**
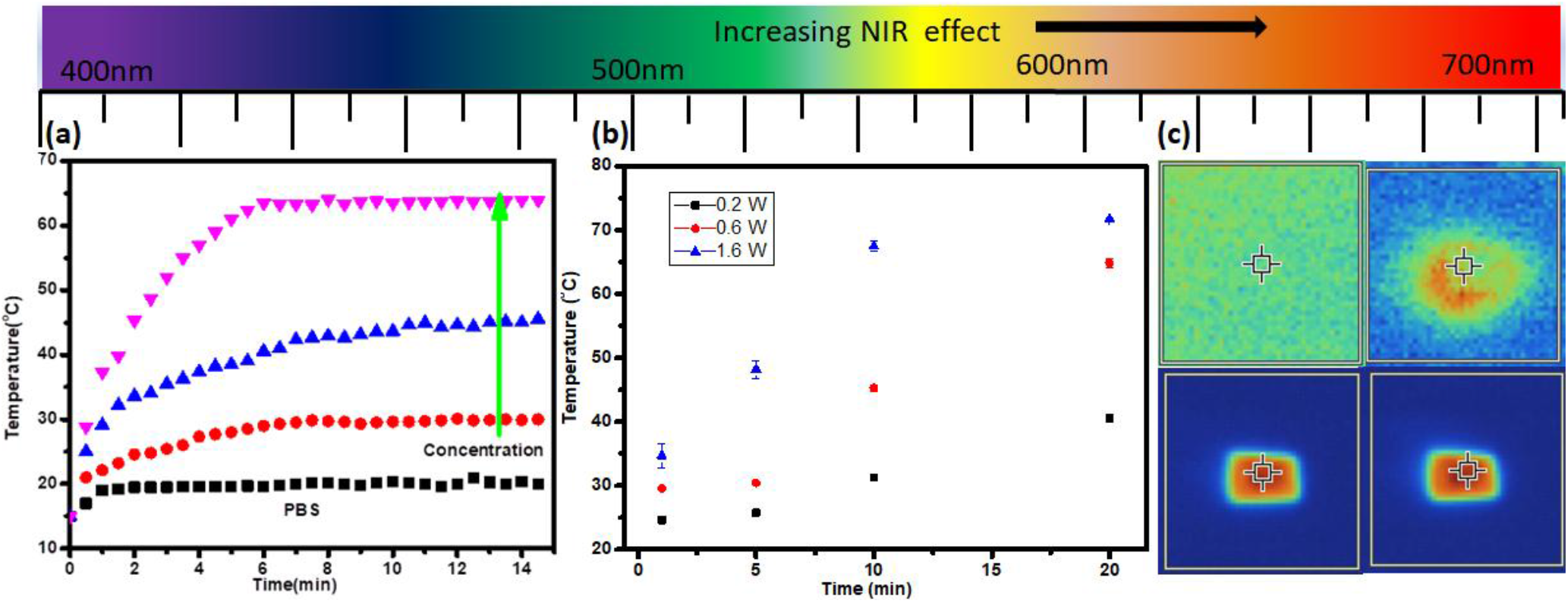
illustration of temperature increases of Film upon NIR **a)** temperature increase PBS vs GNRs solution **b)** laser power depended temperature increase 0.2<0.6<1.6W/cm^-2^ **c)** Thermal elevation in GNR coated thin LBL vs. no GNRs coated thin LBL

Gold nanorods (AuNRs) or nanoparticles (AuNPs) are more chemically stable and biocompatible than silver nanoparticles. Additionally, the near-infrared (NIR) light-induced photothermal effect of gold nanoparticles or nanoparticle aggregates makes them popular for killing bacteria or cancer cells that are resistant to multiple drugs due to their low systemic toxicity, deep penetration, minimal side effects, and spatiotemporally controlled application to the wound.

The PTT effect on GNRs in Gram positive bacteria was studied using the Kirby-Bauer disk diffusion method. Initially the *S*.*aureus* with the concentration of 3×10^8^ CFU/ml was grown on agar plate for 24-48h to maintain the colony stock. To start susceptibility experiment the colony wa transferred on MH agar and prepared LBL was placed onto the inoculate agar plate and irradiated with laser power 1.6 W/cm for 1, 5,10min synergistic killing shown in **Figure (6)** with increasing time and NIR power the zone of inhibition increased.

**Figure 6.**
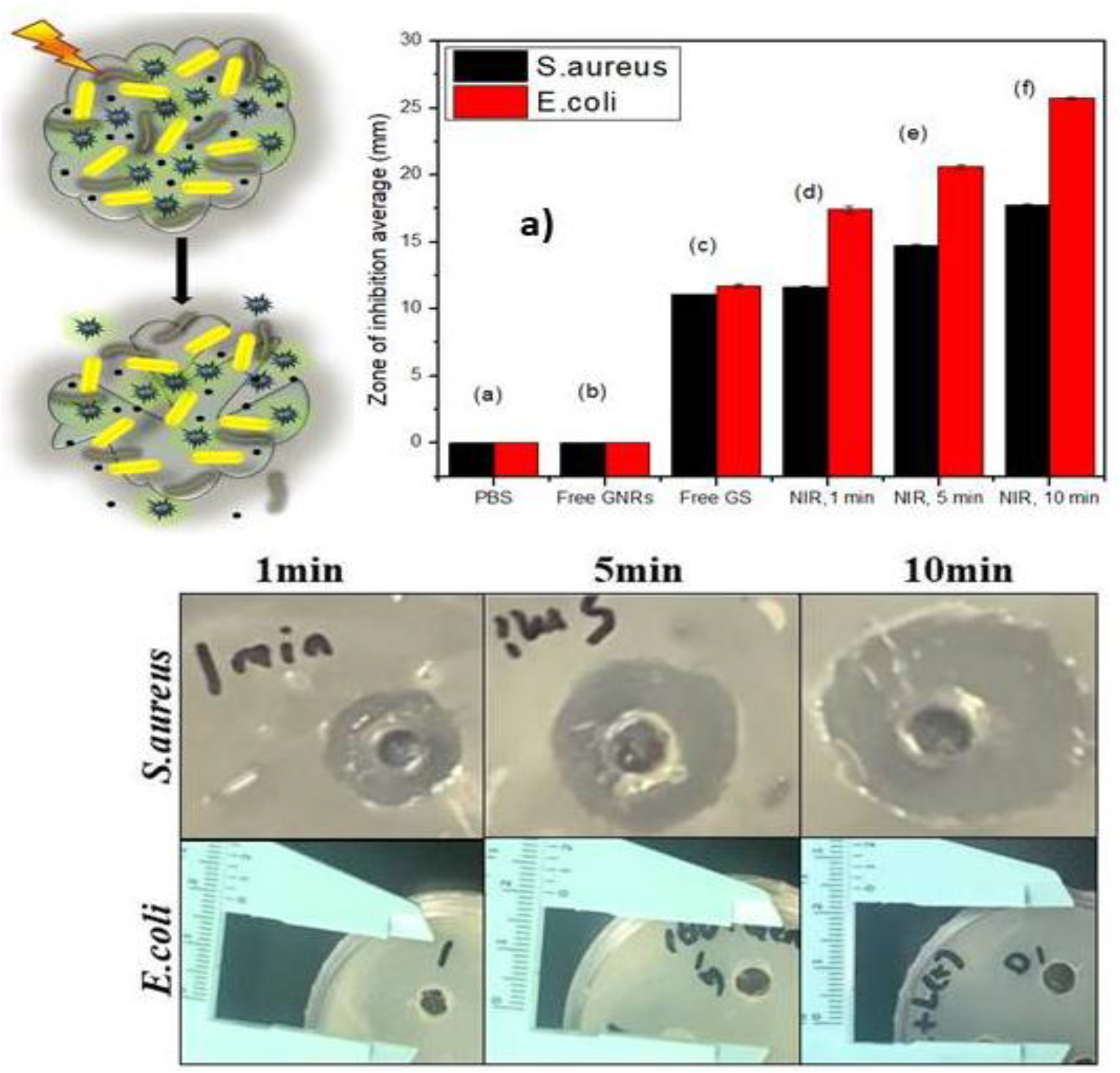
**a)** Zone of inhibition in *S. aureus* and *E*.*coli* depending on NIR + and NIR-using GNRs scaffold, only PBS, FreeGNRs, Free GS, c),d),e),f) PPGSGNRs and NIR+ **b)** image

The bacterial film in chronic wound disturbs the normal wound healing process, this situation demands overdosing of antibiotics within short period, thereby resulting into antibiotic resistance, this limits the therapeutic effects of many pharmaceutical molecules. Moreover, the majority of bacteria live in biofilms, which are composed of organized and coordinated Extracellular polymer compounds safeguard a population of sessile cells (EPS), enabling them to withstand conventional antibiotics and even antibacterial Using metabolic dormancy, nanoparticles can prevent the action of antimicrobial substances via penetration. Chronic and recurrent infections brought on by biofilms enormous financial loads on patients and very high mortality rate societies. The need for efficient and secure antibacterial agents is therefore critical. With fewer opportunities for resistance to emerge to counter the danger from MDR bacterium (multidrug resistant). Furthermore, the antimicrobial substances are also the bacteria were anticipated to penetrate the developed biofilms and be destroyed the biofilm.

## Conclusion

To change the etiology of chronic wound in the presence of drug resistant pathogen is challenging, thus much research is focused to develop ultra-smart nano scaffold considered as smart wound dressing/patch which can provide multiple functionalities. So, in this work we demonstrated fast and simple method to fabricate robust nanoplatform comprised of multilayered polyelectrolyte, assembled by alternate stacking based on electrostatic force, which facilitated loading of multiple drugs. The dense charge on LPEI and PSS help to fabricate multilayer in a well packed manner. Which generated porous pockets thereby helped to embed the pre-functionalized plasmonic nanoparticle and managed to maintain the stability and circulation time? Further to characterize the successfully formed multilayer various techniques were used such as Film thickness (using profilometer), roughness (using AFM), hydrophobicity and hydrophobicity (using contact angle). The dissociation or burst release of pain killer of IBU was obtained in defined pH (7.4) whereas it can be controlled using under different pH range, because in chronic wound exudate there is wide pH range. While assembly the number of layers were importantly considered in order to easily peel off when required, as over layering cab hinder the release of drug at required time was avoided by optimizing the different parameter.

Once the topmost layer dissociated, the next layer with GSGNR became detectable via NIR and followed controlled release of GS which displayed effective results in MIC (gram negative and Gram positive) pathogens. It is noteworthy that this designed nanoscaffold hold good biocompatibility towards normal tissues in wound and is able to response multiple stimuli. Our future work relies on addressing the actual death mechanism using combination therapy in gram negative and gram positive pathogens.

## Supporting information

file:///C:/Users/Nutan/Desktop/LBL%20SPL%2029052026.pdf

