## Supplementary material for "Electrostatically Assembled Layer-by-Layer Plasmonic Implant for Combination Therapy against Multidrug-Resistant Chronic Wound Biofilms": file:///C:/Users/Nutan/Desktop/LBL%20SPL%2029052026.pdf

### Supporting information(S)

**(S1)** Checking loading of model drug Nile red in PPGNRs using different parameter (No. of bilayer, dipping time)

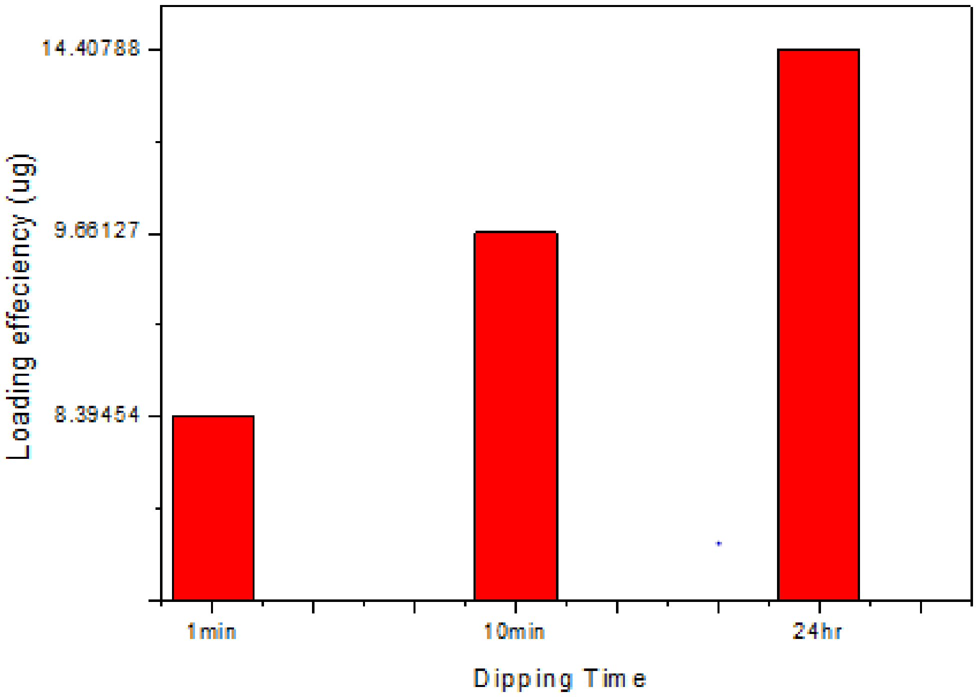


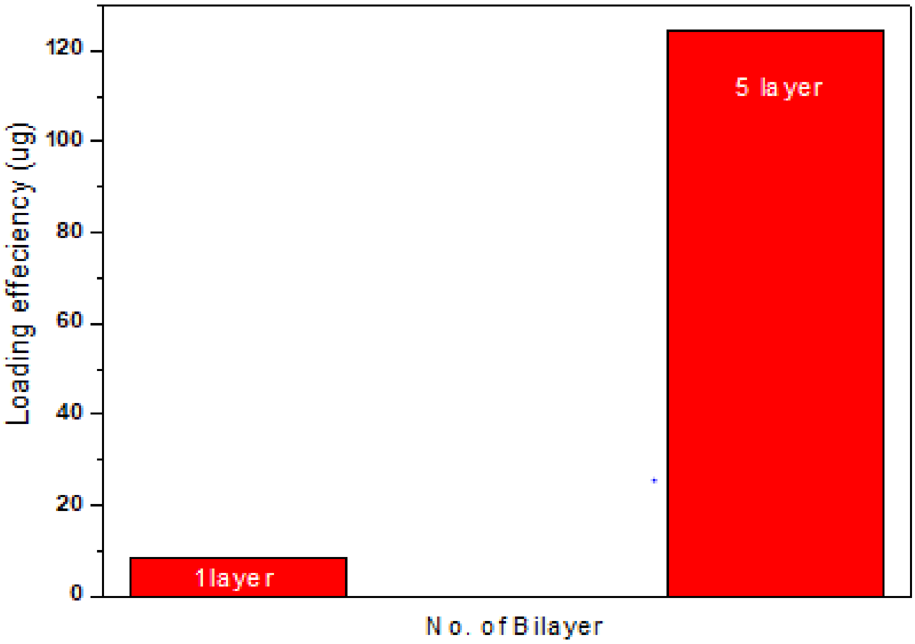

**(S2)** effect of pH in forming thin film

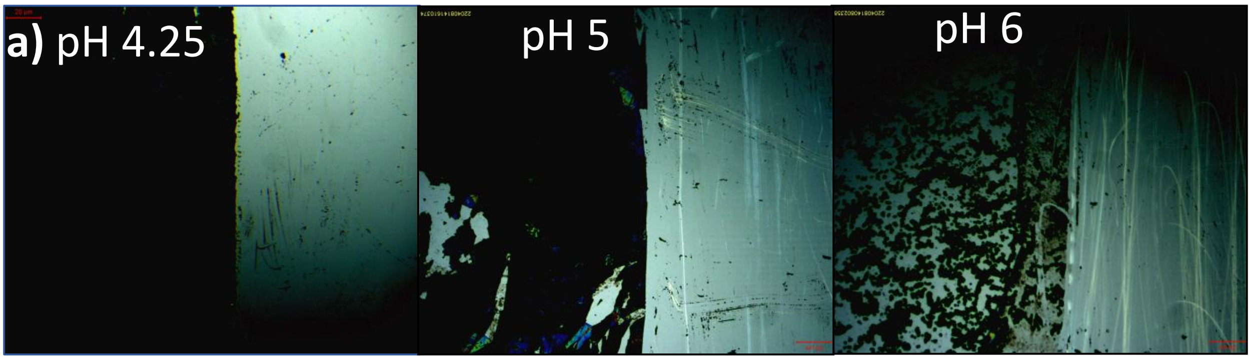


**(S3)** Complete release of IBU for 250 min

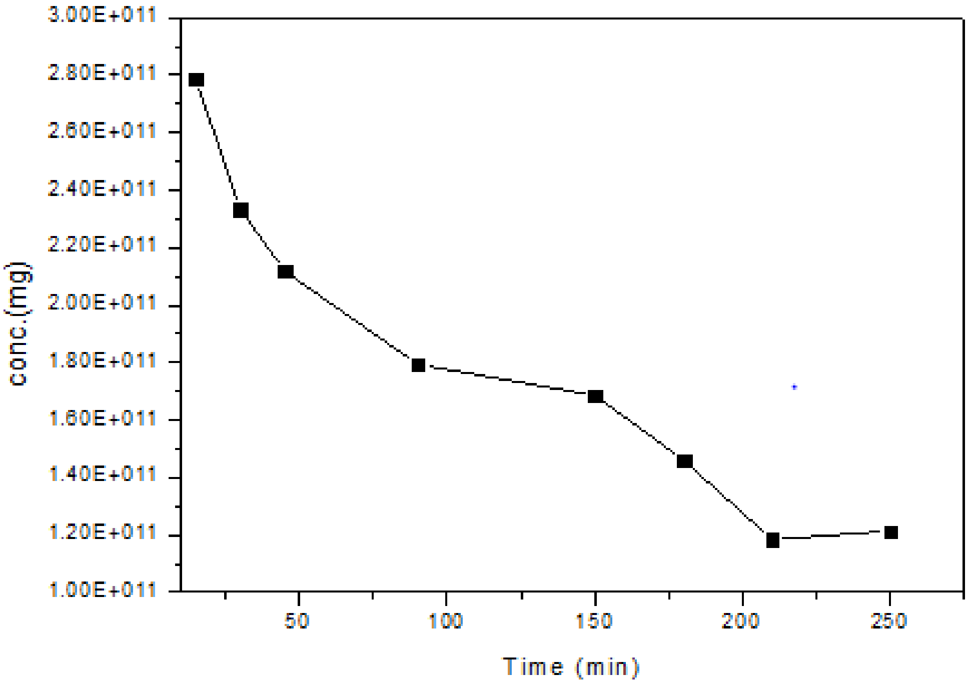
